# Does time matter in phylogeny? A perspective from the fossil record

**DOI:** 10.1101/2021.06.11.445746

**Authors:** Pauline Guenser, Rachel C.M. Warnock, Walker Pett, Philip C.J. Donoghue, Emilia Jarochowska

**Affiliations:** GeoZentrum Nordbayern, Fachgruppe Paläouwelt, University of Erlangen-Nürnberg, Loewenichstr. 28, 91054 Erlangen, Germany; Department of Ecology, Evolution and Organismal Biology, Iowa State University, Ames, IA 50011, USA; School of Earth Sciences, University of Bristol, Life Sciences Building, Tyndall Avenue, Bristol, BS8 1TH UK

## Abstract

The role of time (i.e. taxa ages) in phylogeny has been a source of intense debate within palaeontology for decades and has not yet been resolved fully. The fossilised birth-death range process is a model that explicitly accounts for information about species through time. It presents a fresh opportunity to examine the role of stratigraphic data in phylogenetic inference of fossil taxa. Here, we apply this model in a Bayesian framework to an exemplar dataset of well-dated conodonts from the Late Devonian. We compare the results to those obtained using traditional unconstrained tree inference. We show that the combined analysis of morphology and stratigraphic data under the FBD range process reduces overall phylogenetic uncertainty, compared to unconstrained tree inference. We find that previous phylogenetic hypotheses based on parsimony and stratophenetics are closer to trees generated under the FBD range process. However, the results also highlight that irrespective of the inclusion of age data, a large amount of topological uncertainty will remain. Bayesian inference provides the most intuitive way to represent the uncertainty inherent in fossil datasets and new flexible models increase opportunities to refine hypotheses in palaeobiology.

## INTRODUCTION

Establishing relationships between taxa is important for understanding the evolution and diversification of clades. The rise of model-based inference in morphological phylogenetics has increased the potential for fossils to inform phylogenetic hypotheses [e.g. 1–3]. However, an outstanding question remains about the role of time (i.e. taxon ages) in improving topological accuracy. This stratophenetic *vs*. cladistics debate was intensely pursued in previous decades [4–7]. Some argue that temporal evidence provides a valuable source of evidence for distinguishing between competing phylogenetic hypotheses [8–13] or for testing different phylogenetic methods [14,15]. Others have argued that incompleteness of the fossil record hinders the reliability of stratophenetics or stratolikelihood, and that morphology alone provides a less biased source of evidence for phylogenetic reconstruction [4].

Bayesian inference is becoming increasingly popular in inferring phylogenies in palaeobiology. New models allow us to directly incorporate information about time into phylogenetic inference. The fossilised birth-death (FBD) process unifies the diversification and fossil sampling processes [16] and can be combined with models of character evolution to infer time-calibrated trees [17]. Perhaps unsurprisingly, trees generated under the FBD model show a better fit with the stratigraphic record (i.e. have higher stratigraphic congruence) than unconstrained tree models [18]. However, the impact of fossil ages on topological accuracy is less clear. Simulations suggest that trees generated under the FBD model can be more accurate than unconstrained trees [19,20], but only for datasets that exceed the size of those typical for many fossil groups [20].

Here, we explore the impact of stratigraphic ranges on topology using a new model, the FBD range process [21], which in contrast to the specimen-level FBD process, incorporates information about species durations through time. We chose conodonts as an exemplar to test the impact of stratigraphy on topology using the FBD range model. This group of early vertebrates are characterised by high taxonomic diversity, a sub-continuous fossil record and high abundance, making them one of the most reliable tools for biostratigraphic dating. At the same time, their phylogenetic affinities are highly uncertain, due to the limited availability of morphological characters, which is typical of many fossil datasets. We focus on the late Frasnian-Famennian (Late Devonian) interval, where conodont ranges are especially well constrained.

## MATERIAL AND METHODS

We analysed a matrix of 36 morphological characters based on 15 apparatuses of *Palmatolepis* spp. from Donoghue [9]. The taxa are dated from late Frasnian (*Palmatolepis feisti* biozone) to late Fammenian (*Protognathodus kockeli* biozone). Stratigraphic ranges were obtained from the literature [22–25]. Absolute ages of first and last occurrences were extrapolated from Becker *et al*. [26]. Chronostratigraphic ranges of the species are available in the Supplementary Information (Figure S1).

To compare our results to previous phylogenetic hypotheses, we extracted Helms and Ziegler’ s stratophenetic hypothesis [27], as well as the strict consensus tree under maximum parsimony from Donoghue (TEXT-FIG. 6C and TEXT-FIG. 6A, respectively) [9]. Both trees are available as Supplementary Information (Figure S2, S3).

### Bayesian analyses

Bayesian phylogenetic inference was performed using the software RevBayes [28]. We first analysed the data using traditional (unconstrained) Bayesian tree inference, in the absence of any fossil age information and without a clock model. We used a uniform tree prior on topology and an exponential prior on the branch lengths. The mean of the branch length prior was specified using an exponential hyperprior with rate parameter = 1. The unconstrained tree was rooted using the same outgroup taxa used in previous analyses [9].

We used the Lewis Mk model of morphological character evolution with 3 states, assuming equal rates of change among states [1]. Among character rate variation was modelled using a discretised Gamma distribution with four rate categories, with an exponential prior (rate parameter = 1) on parameter α [29]. Some characters in our dataset are parsimony non-informative, so we specified “ coding = informative” to account for this ascertainment bias [30].

We then analysed the combined morphological and stratigraphic data using the FBD range process as a prior on the tree topology and divergence times [16,21]. We applied a uniform prior to the origin time parameter, constrained between 373.44 Ma (first occurrence of *P. muelleri*) and 378.90 Ma (the base of the Frasnian [26]). We used exponential priors on the speciation, extinction, and fossil recovery rates (mean = 0.1). The probability of sampling at the end of the sampling period (equivalent to extant species sampling parameter ρ) was set to ∼ 0 (0.001, as the program currently does not accept a null value). We assumed a strict clock model (i.e. the rate of discrete character change is assumed to be constant over time), with an exponential prior on the average rate (mean = 1). We used the same substitution model as above.

To explore the impact of excluding sampling ancestors, we removed the sampled ancestor MCMC operators.

For all Bayesian analyses we ran two independent MCMC chains each for a total of 50,000 generations, sampling every 10 generations, resulting in a posterior of 5,000 samples. Maximum clade credibility MCC trees were used to summarise the posterior distribution. To calculate the probability that a given taxa is a sampled ancestor, we used the *SampledAncestorTreeAnalyser* BEAUti application (part of the BEAST2 software package) [17,31].

### Quantitative comparison of the topologies

To quantify the topological differences between the resulting phylogenetic hypotheses, we computed the Robinson-Foulds (RF) distance [32] between consensus trees and all posterior trees for each Bayesian analyses using the *RF*.*dist* function from the *Phangorn* R package [33]. Cladograms from previous studies [9] were transformed into fully binary trees using the *multi2di* function from the *ape* R package [34].

We used RF distances as a distance metric to perform a Non-metric MultiDimensional Scaling (NMDS) analysis using the *metaMDS* function from the *vegan* R package [35]. We randomly subsampled 100 posterior trees from each Bayesian analysis. We computed RF distances between all sets of posterior trees, in addition to the parsimony and stratophenetic trees, resulting in 302 pairwise comparisons. The number of randomly sampled posterior trees were chosen to facilitate convergence of the NMDS analysis. A number of six dimensions was used to reach convergence (stress = 0.10). Although analysis with fewer dimensions did not converge, the graphical patterns we recovered were the same.

Graphical representations of the consensus trees were produced with RStudio [36], using R base and the *RevGadgets* package [37]. Dimensions of the NMDS analysis were plotted with R base and the *viridis* package [38].

## RESULTS

The RF distances show that there is a large discrepancy between the Bayesian MCC trees and the posterior distributions of trees for the same analyses (See Supplementary Information; Figure S4). This indicates that there is a large degree of uncertainty in the overall topology and that the MCC trees are not reliably representative of the posterior distribution. Distance between MCC trees and the posteriors for the same analyses is greater for non-FBD trees (median RF distance = 23 vs. 18; see Supplementary Information; Figure S4). Previous consensus trees, based on parsimony and stratophenetics, are closer to the posterior FBD trees than to the posterior non-FBD trees (median RF distance = 18 - 20 vs. 23 – 25; see Supplementary Information; Figure S4).

The first two dimensions of NMDS analysis, which compares the overall posterior from different Bayesian analyses, also highlights the higher degree of uncertainty associated with the non-FBD output (Figure 1). The FBD trees occupy a more restricted region of the NMDS space than non-FBD trees. This pattern is consistent through the six dimensions of the NMDS analysis (see Supplementary Information; Figure S5). The exclusion of sampled ancestors does not impact the range of uncertainty. The location of the parsimony and stratophenetic trees with respect to the Bayesian trees varies across NMDS dimensions.

**Figure 1:**
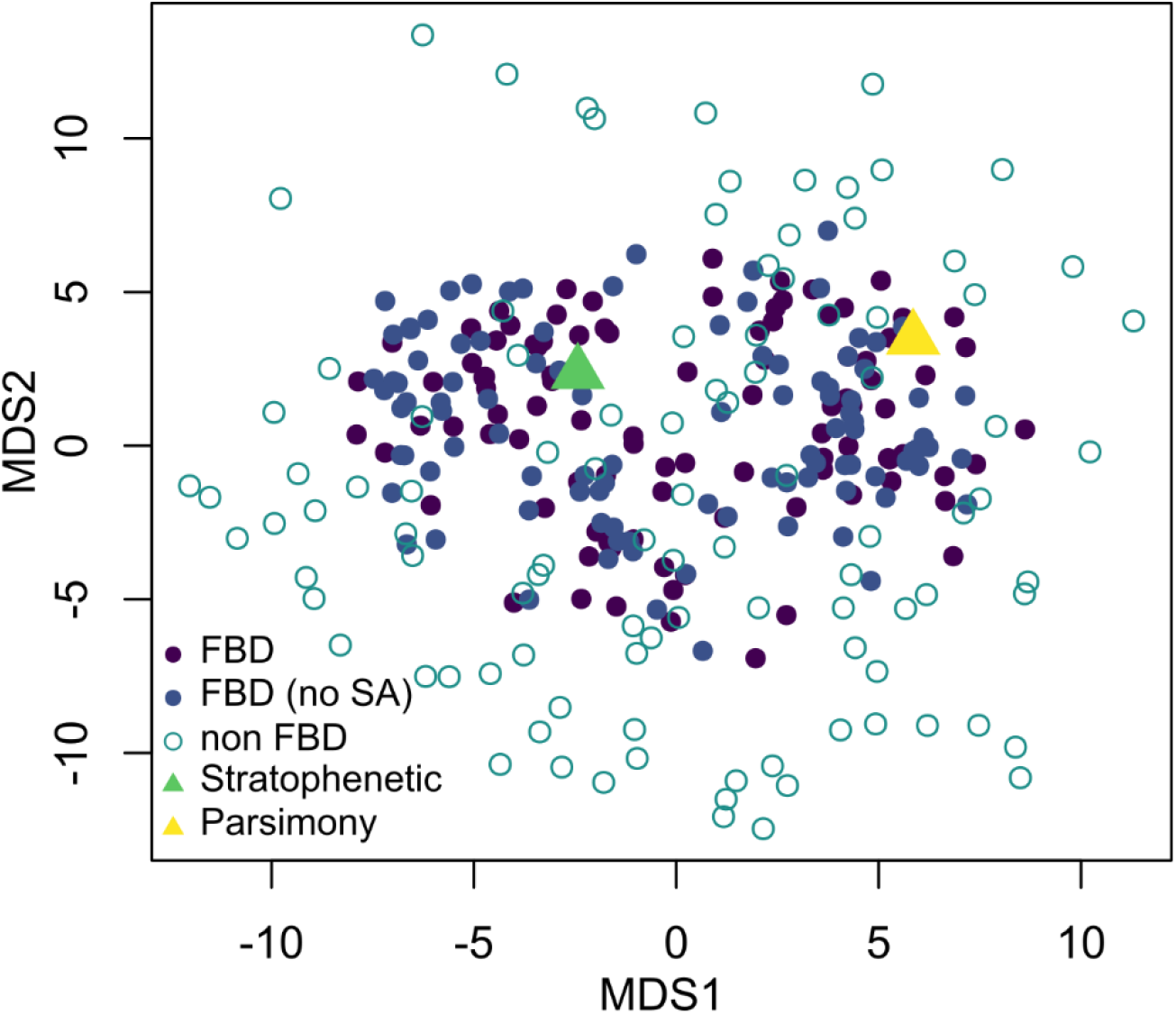
First two dimensions of the NMDS analysis; stress=0.10. The complete pair-wise comparison between the six dimensions are available as supplementary information. Explanations: **FBD** – FBD analysis with sampled ancestors; **FBD (no SA)** – FBD analysis without sampled ancestors; **non FBD** – unconstrained Bayesian analysis; **Stratophenetic** – Helms and Ziegler hypothesis [27], constrained tree from Donoghue [9]; **Parsimony** – Strict consensus tree from Donoghue (TEXT-FIG. 6A) [9].

Despite uncertainty in posterior trees, the clade comprising *P. delicatula delicatula, P. wolskae, P. rugosa, P. schindewolfi, P. perlobata perlobata*, and *P. perlobata maxima* is consistently recovered across analyses (Figure 2). This group is also recovered in the parsimony-based tree, although only a subset (*P. rugosa, P. schindewolfi, P. perlobata perlobata*, and *P. perlobata maxima*, the *P. rugosa* clade) are present in the stratophenetic hypotheses. All other relationships are associated with a high level of uncertainty.

**Figure 2:**
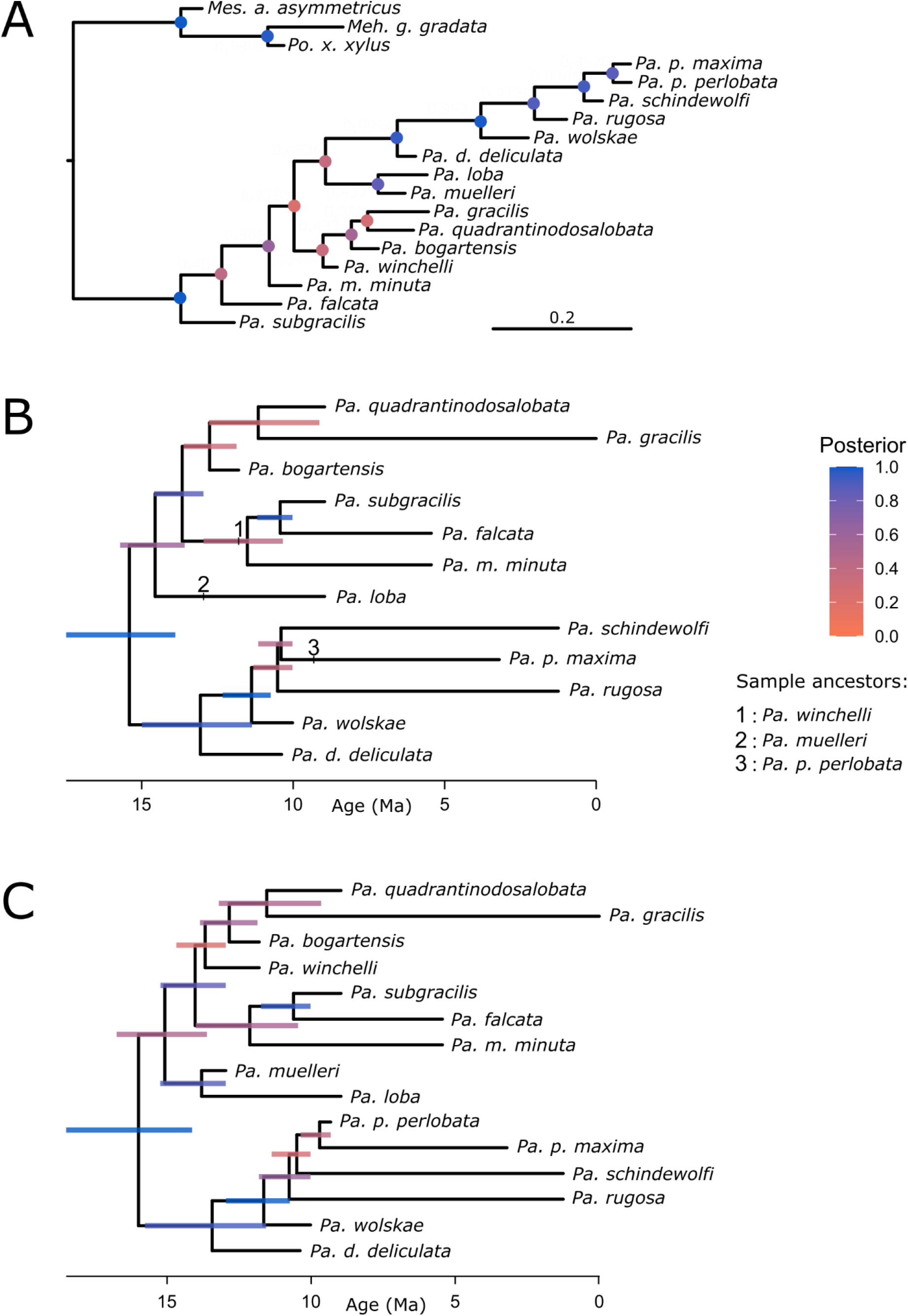
**A** – MCC tree from the unconstrained Bayesian analysis. **B** – MCC tree from FBD range model analysis with sampled ancestors. **C** – MCC tree from FBD range model analysis without sampled ancestors.

The median number of sampled ancestors inferred under the standard FBD model is 3.5. *P. muelleri* and *P. p. perlobata* are strongly supported as sampled ancestors, found in 95.5% and 91.4% of posterior FBD trees, respectively. *P. winchelli*, the third sampled ancestor visible on the FBD MCC tree, is found in 78.7% posterior FBD trees. These results echo the phylogenetic position of these three taxa in the parsimony tree: *P. muelleri* and *P. p. perlobata* are inferred to be sampled ancestors of their respective sister taxa found using parsimony. *P. winchelli* is inferred to be a sampled ancestor of the clade containing *P. subgracilis, P. falcata*, and *P. minuta* (the *P. subgracilis* clade), although this clade is not recovered in the parsimony tree, where *P. winchelli, P. bogartensis, P. quandrantinodosalobata*, and *P. gracilis* (*P. bogartensis* clade) are grouped together.

## DISCUSSION

The recent rise of Bayesian phylogenetic inference in palaeobiology has created the opportunity to revisit the “stratolikelihood” approach to tree building [39,40]. In particular, the fossilised birth-death range process allows for the joint analysis of stratigraphic and morphological data in a statistically coherent framework [21]. Here, we aimed to explore the impact of time (i.e. biostratigraphic ranges) on topology, taking advantage of Late Devonian conodonts that are densely sampled and have well established stratigraphic ranges for this interval.

Our analysis of the *Palmatolepis* morphological character matrix under different probabilistic models (unconstrained Bayesian analysis, and time-constrained analysis using the FBD range model with and without sampled ancestors) shows that the output is associated with substantial uncertainty. This echoes the findings of previous work based on simulated and empirical data showing that small morphological matrices, characteristic of many fossil groups, will not produce precise estimates of topology [15,20,41]. Instead, the pool of trees captured by the posterior, which cannot easily be summarised by a single tree, should be considered the result.

Some authors have argued that stratigraphic ranges can help to overcome the conflict or uncertainty created by morphological convergence [8–13,18]. Fit to the stratigraphic record (i.e. stratigraphic congruence) has therefore been used to measure phylogenetic accuracy [8,42] or compare tree-building methods [14,15,18]. We find that within a Bayesian framework, trees that explicitly incorporate stratigraphic ranges are indeed associated with lower overall uncertainty. This can be seen from the multidimensional representation of RF distances between posterior trees obtained using different models. The NMDS results show that the FBD trees occupy a smaller area of tree space than those occupied by unconstrained trees. This shows that the addition of stratigraphic range data can effectively help to constrain posterior phylogenetic hypotheses.

Trees constructed using parsimony and stratophenetics (i.e. manually constructed) are also on average closer to FBD than unconstrained trees. The disadvantage with non-Bayesian approaches to tree-building is that they do not adequately represent uncertainty. Although our results support the use of time in improving phylogenetic relationships among *Palmatolepis*, it is important to note that the FBD trees are still highly uncertain, which is expected based on the size of our character matrix. However, despite the level of uncertainty, we find persistent support for the *P. rugosa* clade across all methods (parsimony, stratophenetics and Bayesian inference). In addition, we find strong support for the existence of ancestor/descendant relationships – *P. muelleri* and *P. p. perlobata* appear as sampled ancestors in >90% of trees – challenging previous perceptions of *Palmatolepis* spp. relationships.

Previous objections to the use of stratigraphy in phylogenetic inference were that available models could not sufficiently capture the complexity of morphological evolution and that incompleteness of the fossil record made the use of fossil age data unreliable [4,6]. Although phylogenetic models cannot capture the full complexity of evolution, Bayesian inference provides an intuitive way to account for and represent the uncertainty in fossil datasets. Furthermore, as we demonstrate, the inclusion of stratigraphic evidence can be used to reduce the posterior uncertainty in Bayesian phylogenetic inference using the FBD range model. However, future research using simulations are required to examine the impact of stratigraphic incompleteness. We anticipate that extensions of the FBD range model that can better account for the structure of the stratigraphic record will lead to further improvements of phylogenetic inference in deep time.

## DATA AVAILABILITY

All input data, including the stratophenetic and cladistics tree, the matrix of morphological characters and taxon ages are available as Supplementary Information.

## REFERENCES

1. Lewis PO. 2001 A Likelihood Approach to Estimating Phylogeny from Discrete Morphological Character Data. Systematic Biology 50, 913–925. (doi:10.1080/106351501753462876)

2. Nylander JAA, Ronquist F, Huelsenbeck JP, Nieves-Aldrey J. 2004 Bayesian Phylogenetic Analysis of Combined Data. Systematic Biology 53, 47–67. (doi:10.1080/10635150490264699)

3. Wright AM, Lloyd GT, Hillis DM. 2016 Modeling Character Change Heterogeneity in Phylogenetic Analyses of Morphology through the Use of Priors. Systematic Biology 65, 602–611. (doi:10.1093/sysbio/syv122)

4. Smith A. 2000 Stratigraphy in phylogeny reconstruction. Journal of Paleontology 74, 763–766. (doi:10.1666/0022-3360(2000)074<0763:SIPR>2.0.CO;2)

5. Wagner PJ. 2002 Testing phylogenetic hypothese with stratigraphy and morphology - A comment on Smith (2000). Journal of Paleontology 76, 590–593. (doi:10.1666/0022-3360(2002)076<0590:TPHWSA>2.0.CO;2)

6. Smith A. 2002 Stratigraphy in phylogeny reconstruction - Respons. Journal of Paleontology 76, 594–595. (doi:10.1666/0022-3360(2002)076<0594:SIPRR>2.0.CO;2)

7. Fisher DC, Foote M, Fox DL, Leighton L. 2002 Stratigraphy in phylogeny reconstruction - Comment on Smith (2000). Journal of Paleontology 76, 585–586. (doi:10.1666/0022-3360(2002)076<0585:SIPRCO>2.0.CO;2)

8. Wagner PJ. 1995 Stratigraphic Tests of Cladistic Hypotheses. Paleobiology 21, 153–178.

9. Donoghue PCJ. 2001 Conodonts Meet Cladistics: Recovering Relationships and Assessing the Completeness of the Conodont Fossil Record. Palaeontology 44, 65–93. (doi:10.1111/1475-4983.00170)

10. Dzik J. 1991 Features of the fossil record of evolution. Acta Palaeontologica Polonica 36.

11. Dzik J. 1999 Relationship between rates of speciation and phyletic evolution: Stratophenetic data on pelagic conodont chordates and benthic ostracods. Geobios 32, 205–221. (doi:10.1016/S0016-6995(99)80033-3)

12. Steineck PL, Fleisher RL. 1978 Towards the Classical Evolutionary Reclassification of Cenozoic Globigerinacea (Foraminiferida). Journal of Paleontology 52, 618–635.

13. Lazarus DB, Prothero DR. 1984 The Role of Stratigraphic and Morphologic Data in Phylogeny. Journal of Paleontology 58, 163–172.

14. Sansom RS, Choate PG, Keating JN, Randle E. 2018 Parsimony, not Bayesian analysis, recovers more stratigraphically congruent phylogenetic trees. Biology Letters 14, 20180263. (doi:10.1098/rsbl.2018.0263)

15. Wright AM, Lloyd GT. 2020 Bayesian analyses in phylogenetic palaeontology: interpreting the posterior sample. Palaeontology 63, 997–1006. (doi:10.1111/pala.12500)

16. Stadler T. 2010 Sampling-through-time in birth–death trees. Journal of Theoretical Biology 267, 396–404. (doi:10.1016/j.jtbi.2010.09.010)

17. Gavryushkina A, Heath TA, Ksepka DT, Stadler T, Welch D, Drummond AJ. 2016 Bayesian Total-Evidence Dating Reveals the Recent Crown Radiation of Penguins. Systematic Biology 66, 57–73. (doi:10.1093/sysbio/syw060)

18. King B. 2021 Bayesian Tip-Dated Phylogenetics in Paleontology: Topological Effects and Stratigraphic Fit. Systematic Biology 70, 283–294. (doi:10.1093/sysbio/syaa057)

19. Mongiardino Koch N, Garwood RJ, Parry LA. 2021 Fossils improve phylogenetic analyses of morphological characters. Proceedings of the Royal Society B: Biological Sciences 288, 20210044. (doi:10.1098/rspb.2021.0044)

20. Barido-Sottani J, van Tiel NMA, Hopkins MJ, Wright DF, Stadler T, Warnock RCM. 2020 Ignoring Fossil Age Uncertainty Leads to Inaccurate Topology and Divergence Time Estimates in Time Calibrated Tree Inference. Frontiers in Ecology and Evolution 8, 183. (doi:10.3389/fevo.2020.00183)

21. Stadler T, Gavryushkina A, Warnock RCM, Drummond AJ, Heath TA. 2018 The fossilized birth-death model for the analysis of stratigraphic range data under different speciation modes. Journal of Theoretical Biology 447, 41–55. (doi:10.1016/j.jtbi.2018.03.005)

22. Klapper G. 1988 The Montagne Noire Frasnian (Upper Devonian) Conodont Succession., 449–468.

23. Klapper G. 2007 Conodont taxonomy and the recognition of the Frasnian/Famennian (Upper Devonian) stage boundary. Stratigraphy 4, 67–76.

24. Spalletta C, Perri MC, Over DJ, Corradini C. 2017 Famennian (Upper Devonian) conodont zonation: revised global standard.

25. Ovnatanova NS, Kononova LI. 2020 Taxonomic problems of some zonal species of the genus Palmatolepis and the correlation of the Frasnian of the East European Platform with conodont zonations. 70, 107–124.

26. Becker RT, Marshall JEA, Da Silva A-C, Agterberg FP, Gradstein FM, Ogg JG. 2020 Chapter 22 - The Devonian Period. In Geologic Time Scale 2020 (eds FM Gradstein, JG Ogg, MD Schmitz, GM Ogg), pp. 733–810. Elsevier. (doi:10.1016/B978-0-12-824360-2.00022-X)

27. Moore RC, Robison RA, Geological society of America, editors. 1981 Treatise on invertebrate paleontology Part W Miscellanea Supplement 2 Conodonta. Boulder (Colo.), Lawrence, Kan.: Geological Society of America, University of Kansas.

28. Höhna S, Landis MJ, Heath TA, Boussau B, Lartillot N, Moore BR, Huelsenbeck JP, Ronquist F. 2016 RevBayes: Bayesian phylogenetic inference using graphical models and an interactive model-specification language. Systematic Biology 65, 726–736. (doi:10.1093/sysbio/syw021)

29. Yang Z. 1994 Maximum likelihood phylogenetic estimation from DNA sequences with variable rates over sites: approximate methods. Journal of Molecular evolution 39, 306–314.

30. Allman ES, Rhodes JA. 2008 Identifying evolutionary trees and substitution parameters for the general Markov model with invariable sites. Mathematical Biosciences 211, 18–33. (doi:10.1016/j.mbs.2007.09.001)

31. Bouckaert R et al. 2019 BEAST 2.5: An advanced software platform for Bayesian evolutionary analysis. PLOS Computational Biology 15, e1006650. (doi:10.1371/journal.pcbi.1006650)

32. Robinson DF, Foulds LR. 1981 Comparison of phylogenetic trees. Mathematical Biosciences 53, 131–147. (doi:10.1016/0025-5564(81)90043-2)

33. Schliep KP. 2011 phangorn: phylogenetic analysis in R. Bioinformatics 27, 592–593. (doi:10.1093/bioinformatics/btq706)

34. Paradis E, Schliep K. 2019 ape 5.0: an environment for modern phylogenetics and evolutionary analyses in R. Bioinformatics 35, 526–528. (doi:10.1093/bioinformatics/bty633)

35. Oksanen J et al. 2020 vegan: Community ecology package. R package version 2. 5–7.

36. RStudio Team. 2020 RStudio: Integrated Development for R. PBC, Boston, MA. See http://www.rstudio.com/.

37. Tribble CM, Freyman WA, Landis MJ, Lim JY, Barido-Sottani J, Kopperud BT, Höhna S, May MR. 2021 RevGadgets: an R Package for visualizing Bayesian phylogenetic analyses from RevBayes. bioRxiv, 2021.05.10.443470. (doi:10.1101/2021.05.10.443470)

38. Garnier N, Ross N, Rudis R, Carmargo AP, Sciaini M, Scherer C. 2021 Rvision - Colorblind-Friendly Color Maps for R.

39. Wagner PJ. 2000 Likelihood tests of hypothesized durations: determining and accommodating biasing factors. Paleobiology 26, 431–449. (doi:10.1666/0094-8373(2000)026<0431:LTOHDD>2.0.CO;2)

40. Wagner PJ. 1999 The utility of fossil data in phylogenetic analyses: a likelihood example using Ordovician-Silurian species of the Lophospiridae (Gastropoda: Murchisoniina). American Malacological Bulletin 15, 1–31.

41. O’ Reilly JE, Donoghue PCJ. 2018 The efficacy of consensus tree methods for summarizing phylogenetic relationships from a posterior sample of trees estimated from morphological data. Systematic Biology 67, 354–362. (doi:10.1093/sysbio/syx086)

42. Benton MJ, Hitchin R. 1997 Congruence between phylogenetic and stratigraphic data on the history of life. Proceedings of the Royal Society of London. Series B: Biological Sciences 264, 885–890. (doi:10.1098/rspb.1997.0123)

