## Supporting Information for "Does time matter in phylogeny? A perspective from the fossil record"

### Supplementary Information

Pauline GUENSER, Rachel C.M. WARNOCK, Walker PETT, Philip C.J. DONOGHUE, Emilia JAROCHOWSKA

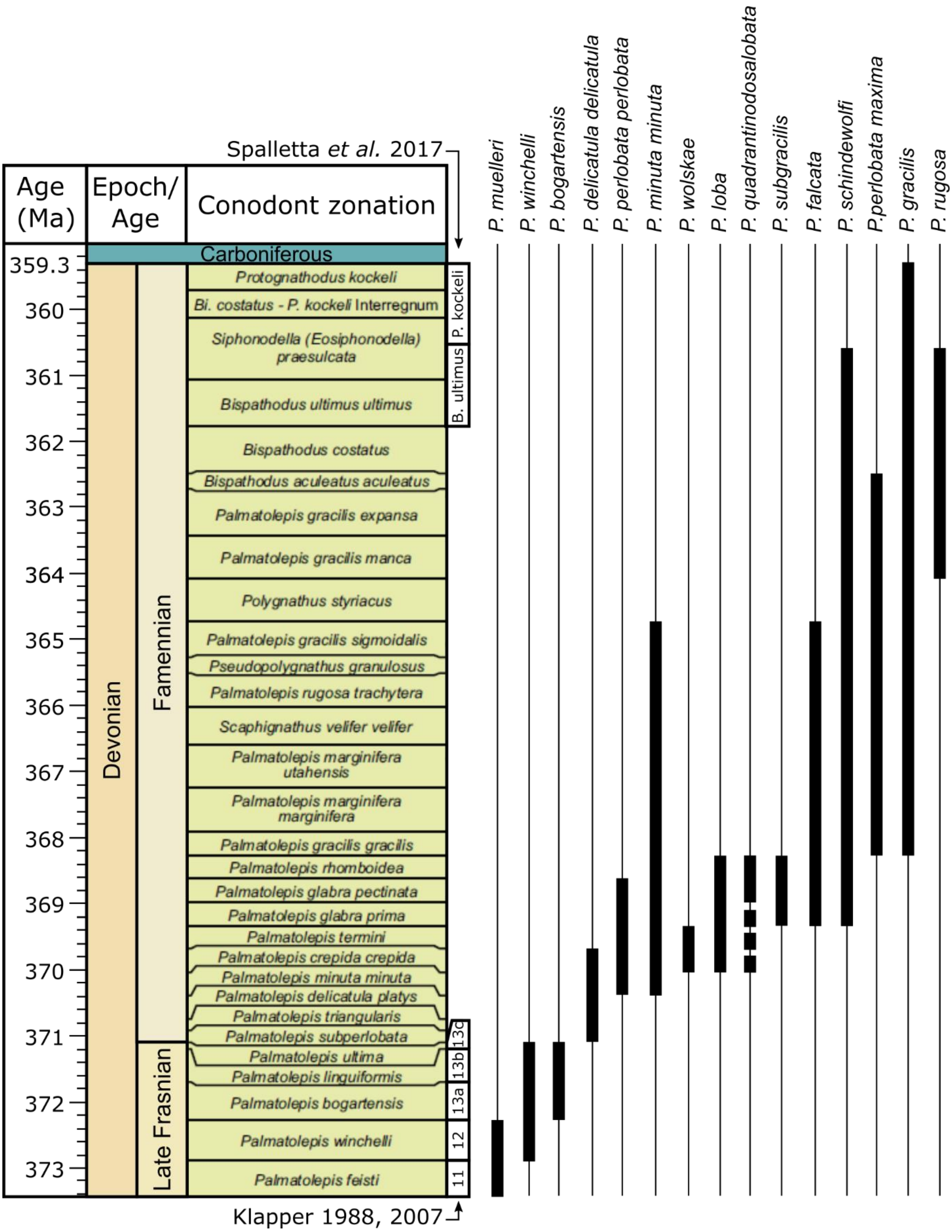


*Figure S 1 : Biostratigraphic ranges of the 15* Palmatolepis *species analysed in this study [1–5]. Chronostratigraphic scheme and biozonation based on Becker* et al. *[2].*

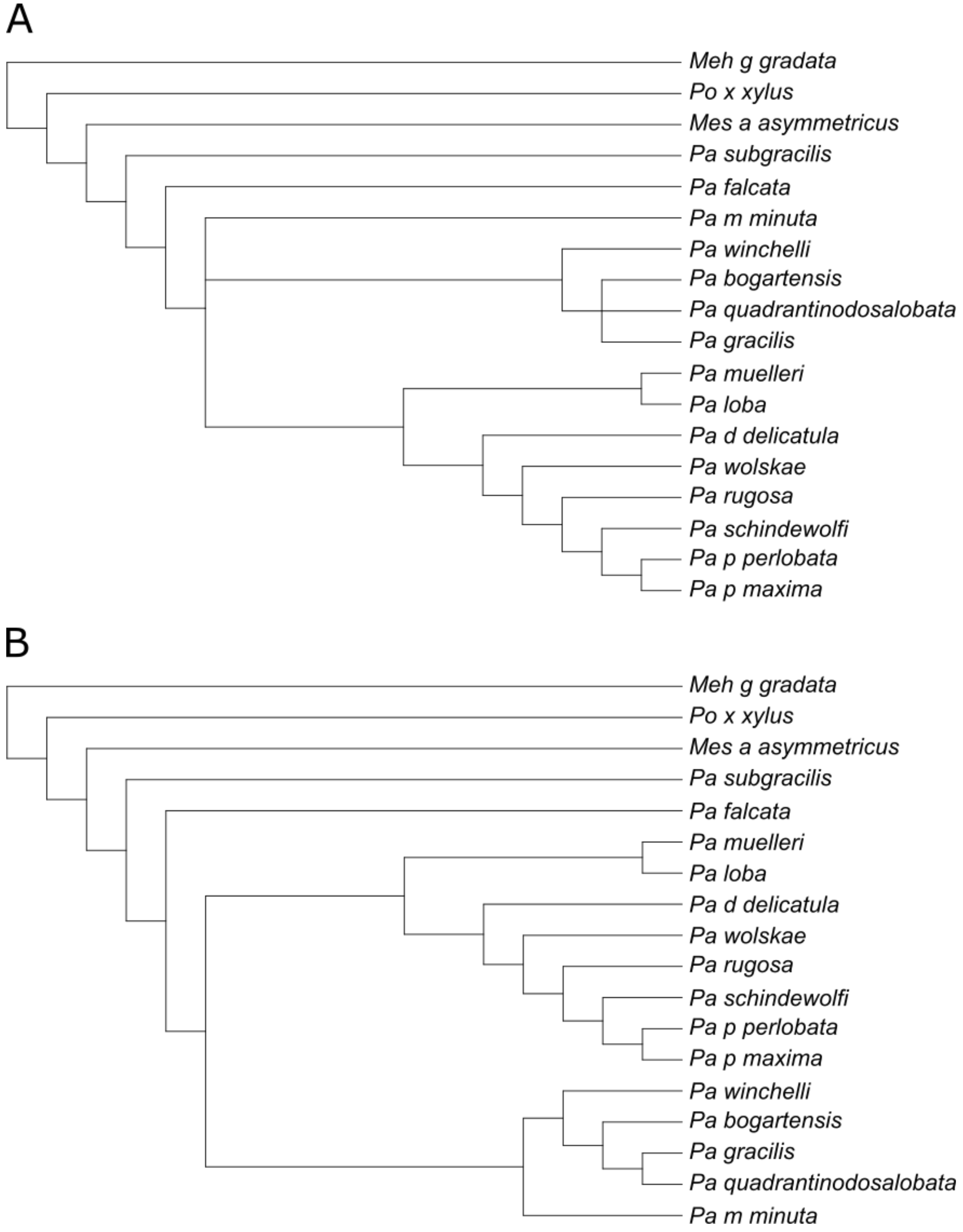


*Figure S 2: Strict consensus tree from Donoghue (TEXT-FIG. 6A) [6].* ***A*** *– Original hypothesis.* ***B*** *– Binary tree using* multi2di *function from* ape *R package.*

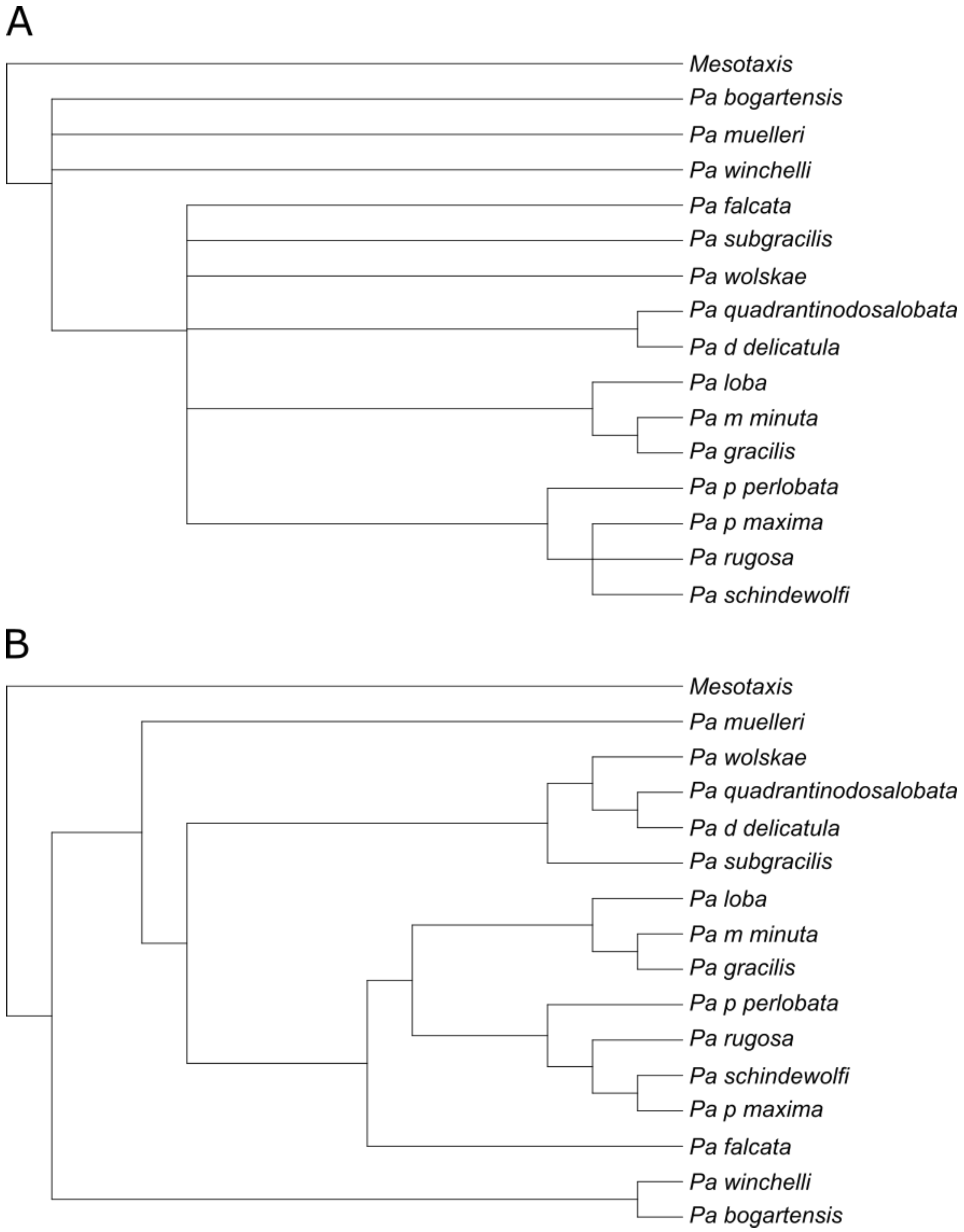


*Figure S 3 : Stratophenetic hypothesis based on Helms qnd Ziegler [7].* ***A*** *– Constrained tree built by Donoghue (TEXT-FIG. 6C) [6].* ***B*** *– Binary tree using* multi2di *function from* ape *R package.*

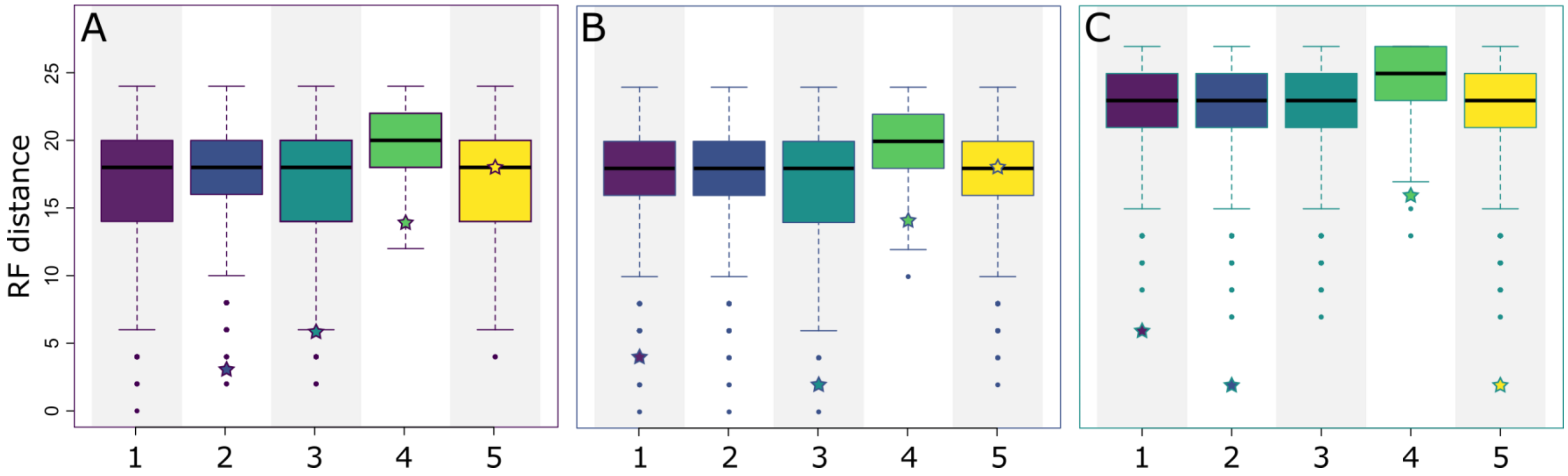


*Figure S 4 : RF distances between posteriors and consensus trees. Stars represent RF distance between consensus trees.* ***A*** *– RF distances between FBD posteriors and consensus trees.* ***B*** *– RF distances between FBD posteriors without sampled ancestors and consensus trees.* ***C*** *– RF distances between unconstrained Bayesian analysis and consensus trees.* ***1*** *(purple boxplot) – FBD MCC tree;* ***2*** *(blue boxplot) – FBD MCC tree without sampled ancestors;* ***3*** *(dark cyan boxplot) – Unconstrained MCC tree;* ***4*** *(green boxplot) – Stratophenetic tree (based on Helms and Ziegler's hypothesis [7], from Donoghue (TEXT-FIG. 6C) [6]);* ***5*** *(yellow boxplot) – Strict consensus tree under maximum parsimony from Donoghue (TEXT-FIG. 6A) [6].*

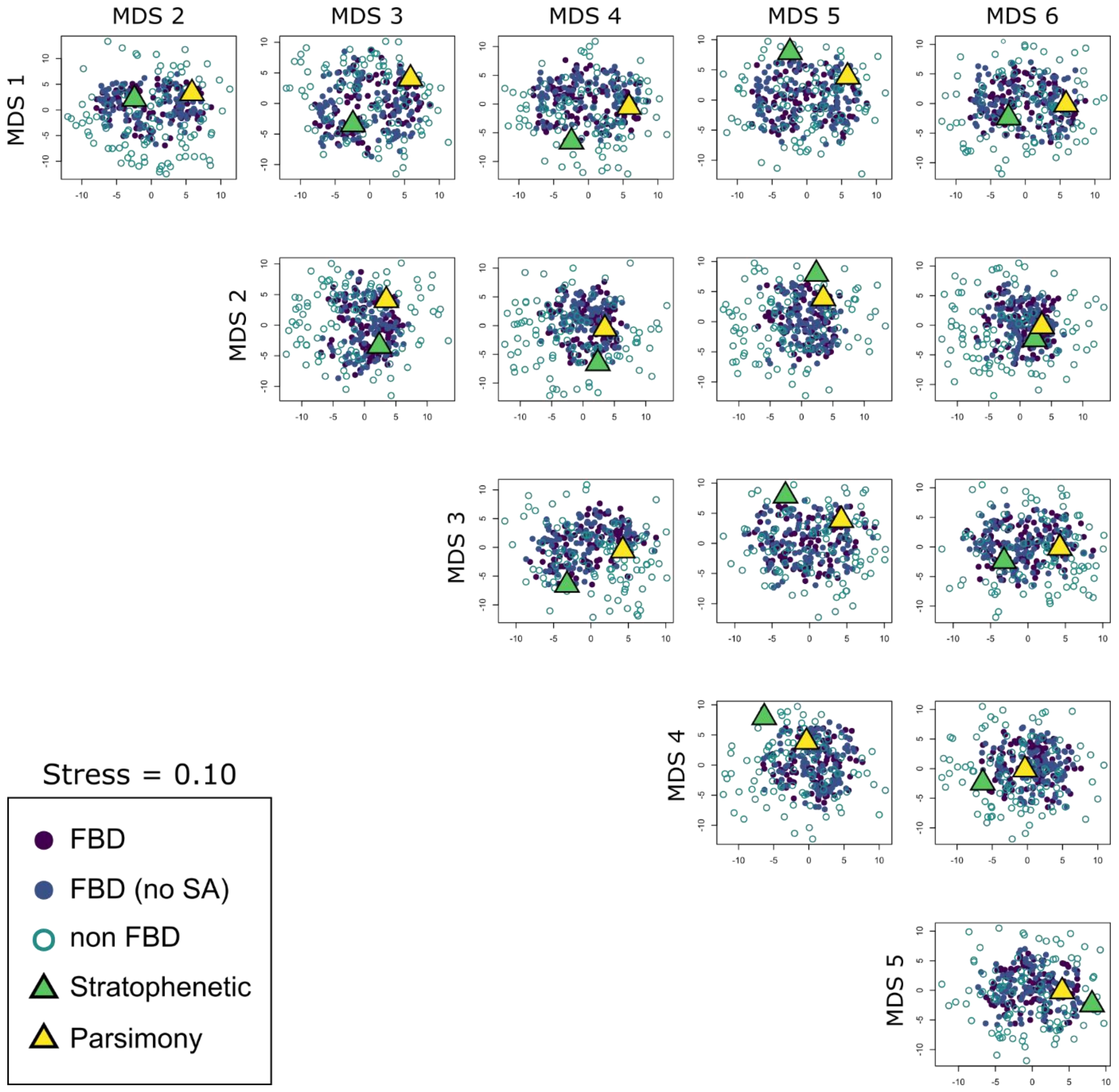


*Figure S 5 : Representation of the six dimensions of the NMDS. Explanations:* ***FBD*** *– FBD analysis with sampled ancestors;* ***FBD (no SA)*** *– FBD analysis without sampled ancestors;* ***non FBD*** *– unconstrained Bayesian analysis;* ***Stratophenetic*** *– Helms and Ziegler hypothesis [7], constrained tree from Donoghue [6];* ***Parsimony*** *– Strict consensus tree from Donoghue (TEXT-FIG. 6A) [6]. The use of* metaMDS vegan *function allows to order the dimensions by their importance in terms of variance explanation. As such, the first dimension holds more information than the second dimension, which holds more information than the third dimension, and so on.*

### REFERENCES


1. Spalletta C, Perri MC, Over DJ, Corradini C. 2017 Famennian (Upper Devonian) conodont zonation: revised global standard.

2. Becker RT, Marshall JEA, Da Silva A-C, Agterberg FP, Gradstein FM, Ogg JG. 2020 Chapter 22 - The Devonian Period. In *Geologic Time Scale 2020* (eds FM Gradstein, JG Ogg, MD Schmitz, GM Ogg), pp. 733–810. Elsevier. (doi:10.1016/B978-0-12-824360-2.00022-X)

3. Klapper G. 1988 The Montagne Noire Frasnian (Upper Devonian) Conodont Succession. , 449–468.

4. Klapper G. 2007 Conodont taxonomy and the recognition of the Frasnian/Famennian (Upper Devonian) stage boundary. *Stratigraphy* **4**, 67–76.

5. Ovnatanova NS, Kononova LI. 2020 Taxonomic problems of some zonal species of the genus Palmatolepis and the correlation of the Frasnian of the East European Platform with conodont zonations. **70**, 107–124.

6. Donoghue PCJ. 2001 Conodonts Meet Cladistics: Recovering Relationships and Assessing the Completeness of the Conodont Fossil Record. *Palaeontology* **44**, 65–93. (doi:10.1111/1475-4983.00170)

7. Moore RC, Robison RA, Geological society of America, editors. 1981 *Treatise on invertebrate paleontology Part W Miscellanea Supplement 2 Conodonta*. Boulder (Colo.), Lawrence, Kan.: Geological Society of America, University of Kansas.
